# BaitBench: a comprehensive tool for building, assessing, and predicting outcomes for hybrid capture enrichment probes

**DOI:** 10.64898/2026.10.24.761557

**Authors:** Aniello M. Infante, Brandon D. Hollingsworth, Shaun T. Cross

**Affiliations:** Department of Environmental, Agricultural, and Occupational Health, University of Nebraska Medical Center, Omaha, Nebraska, USA; Department of Epidemiology and Biostatistics, Arnold School of Public Health, University of South Carolina, Columbia, SC

## Abstract

The effectiveness of hybridization capture sequencing, a powerful lab technique to isolate and characterize low-abundance samples from complex nucleic acid mixtures, relies heavily on creating a robust probe set. There are a variety of tools that design, evaluate, or simulate capture probes, however, these tools are often siloed and frequently difficult to use. To our knowledge there is no tool that connects all steps of the hybridization capture probe development and analysis process. Here we introduce BaitBench, a comprehensive tool to aid in the design, assessment, and simulation of target capture probes. BaitBench wraps or reimplements in Rust several current tools, and implements a thermodynamic model for simulating capture. Integrating steps allows us to track simulated reads through the whole process enabling us to report sources of false positives and reasons for false negatives. Easy to read output files and the ability to run via command line or GUI interfaces increases usability. We validated BaitBench against publicly available data and achieved a high correlation (*ρ* = 0.916), demonstrating its accuracy and translation to real-world settings. Code is available at github.com/niel-infante/BaitBench, with installers for Mac and Windows available at github.com/niel-infante/BaitBench/releases.

## 1 Introduction

Hybridization-based target capture selectively enriches genomic targets from complex nucleic acid mixtures by hybridizing biotinylated oligonucleotide probes to complementary library fragments, pulling down probe-bound material with streptavidin beads, and sequencing the enriched fraction. The result is an orders-of-magnitude increase in on-target read depth, making the approach essential wherever targets represent a small fraction of total nucleic acid. Example uses include clinical metagenomics, pathogen surveillance, antimicrobial resistance gene detection, viral genomics, ancient DNA, and whole-exome sequencing (Bravo et al. 2025). In these settings, where targets may constitute as little as 0.001–1% of the total nucleic acid, probe panel design and performance can be the decisive factors between a successful assay and a failed one.

Tools such as CATCH (Metsky et al. 2019), Syotti (Alanko et al. 2022), and ProbeTools (Kuchinski et al. 2022) can efficiently construct probe sets with guaranteed coverage across diverse, rapidly evolving targets. However, a systematic gap remains between designing a panel and predicting how it will perform. No broadly adopted approach exists for estimating sensitivity as a function of target abundance, specificity against host or background sequences, within-panel cross-reactivity, or coverage uniformity before probes are synthesized and wet-lab validation begins. Wet-lab iteration is expensive, slow, and labor-intensive. In silico evaluation is cheap and fast but underutilized, in part because it requires assembling multiple tools with incompatible formats and dependencies, a major barrier for users without specialized bioinformatics expertise. When simulations are used, they typically assume uniform per-base capture probability, ignoring the sequence-dependent binding affinities that govern real hybridization. Recent work with RAmpSim (Zhang et al. 2025) demonstrated that thermodynamic simulation using the SantaLucia nearest-neighbor model (SantaLucia 1998) produces substantially more realistic hybridization capture, but RAmpSim addresses only the simulation step and does not include probe design, quality control, or quantitative performance metrics.

Here we present BaitBench, an end-to-end suite for designing, assessing, and benchmarking probe panels. BaitBench integrates Rust reimplementations of CATCH and Syotti with a quality-controlled build pipeline, alignment-based probe assessment, and a full thermodynamic simulation of the capture-sequencing workflow enhancing RAmpSim with AT/GC initiation penalties. A three-way classification scheme separately quantifies within-panel cross-reactivity and true off-target capture, a distinction that binary detected/not-detected calls obscure. Written in Rust and distributed with an optional desktop GUI, BaitBench is designed to lower the barrier to rigorous probe panel evaluation for command-line and non-specialist users alike.

## 2 Features and Functionality

### Building Probes

The pipeline starts with quality control (QC) of input sequences by collapsing redundant sequences with cd-hit (Li and Godzik 2006), removing sequences of more than 5% ambiguous bases, and removing short sequences (default to probe length). Options are a simple tiling algorithm, wrapped CATCH, or native Rust reimplementations of CATCH (Metsky et al. 2019) and Syotti (Alanko et al. 2022). With a user flag, any ambiguous nucleotide (N) in a probe will be replaced by a nucleotide based on surrounding sequence, a GC content filter is applied, and then probes with low complexity are removed by an internal implementation of sDust (Morgulis et al. 2006).

### Assessing Probes

The assess-probes module aligns probes to targets using minimap2 (Li 2018) to assess target coverage and probe multimapping. Probes are mapped to themselves to assess cross reactivity, and to background genomes to identify off-target binding and probe specificity. Statistics are compiled and further mapping is done to assess trouble spots. Everything is displayed in an interactive report which will show coverage gaps for targets, give insight into probe performance, identify redundant probes, and provide an analysis of uncovered regions.

### Simulating Capture

Assessing probes assumes a (near) perfect world. To simulate a more realistic capture experiment BaitBench implements an eight-step process (Prepare, Simulate, Sequence, Filter, Map, List, Metrics, Report). Each step can be run separately and all intermediate files are documented and retained; the pipeline can be re-entered at any step.

1. **Prepare** creates a single fasta file containing all sequences, along with a weights file that specifies what is in the simulated sample, and in what proportion. Inputs include targets, target genomes (when the targets are not whole genomes), distractors, and host. Weights are calculated from user weighting, genome size, or distractor fractions.
2. **Simulate** aligns all probes to possible binding locations, Gibbs free energy is calculated, and fragments are generated randomly weighted by thermodynamic properties and sequence input weight.
3. **Sequence** takes the fragments captured and models the actual sequencing step either perfectly via fragment trimming, or using one of the wrapped sequence simulators, ART-modern (Yu 2026) or Badread (Wick 2019).
4. **Filter** is an optional step that maps the simulated reads to the host reference, and only passes on reads that do not map.
5. **Map** aligns reads to the target sequences
6. **List** parses the SAM output and counts reads per reference sequence.
7. **Metrics** computes a three way classification of true/false negative/positive target/distractor hits.
8. **Report** produces a self-contained HTML report generated via RMarkdown. A more detailed discussion of each of these steps is available in the BaitBench documentation.

Simulate is of high importance as it predicts probe efficiency for the hybridization step. Our thermodynamic model follows RAmpSim (Zhang et al. 2025) with the addition of AT/GC initiation penalties. Probes are aligned promiscuously to the combined reference with minimap2; CIGAR and MD tags are parsed to reconstruct per-position base pairs for each alignment. Gibbs free energy is then calculated for each probe–reference duplex using the SantaLucia nearest-neighbor model (SantaLucia 1998):

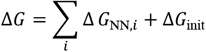

where the sum runs over consecutive nearest-neighbor dinucleotide steps, while Δ*G*_init_ applies AT/GC initiation penalties.

For each probe-binding site pair, ΔG is converted to a Boltz-mann-weighted binding score *w* ∝ *e*^−Δ*G*/*RT*^. Fragment sampling then proceeds in two waves, captured fragments and background fragments. For captured fragments, a probe is drawn uniformly from all probes with at least one alignment. Then, a specific alignment site for that probe is sampled weighted by *w* and the abundance of the target sequence. A fragment is generated with a random center offset and and length drawn from a truncated normal distribution. In the second wave, background fragments are sampled uniformly across the reference weighted by sequence length and abundance. A user-specified capture fraction sets the ratio of probe-biased fragments to background fragments. The relative enrichment of each target emerges from the thermodynamic sampling rather than being imposed as an explicit per-target parameter.

### Sweeping Parameters

In addition to individual simulations, BaitBench allows users to do a sweep over some key parameters: capture fraction, temperature, number of sequences generated, and the initial fraction of desired sample present. The resulting coverage curve gives users translational insights into sequencing depths needed to reach coverage sufficient for their downstream analyses.

As an example, **Figure 1** shows coverage depth curves for Zika virus (ZIKV) detection against an *Aedes aegypti* background using a probe set built against ZIKV sequences. Panels show different capture fractions, and lines indicate number of simulated reads. At a capture fraction of 0.02 (top left), a complete Zika virus genome is impossible to capture regardless of read count. However, if we can slightly increase the capture fraction to 0.05–0.1 (e.g. by increasing hybridization time or improving wash stringency), identification of ZIKV presence is possible even at low read counts, and near-complete genome coverage is acheived with 25,000–50,000 reads. However, if for example we wanted greater insight into well-supported genetic variants (i.e. higher coverage), significant increases in capture fraction and/or total read count are required.

**Figure 1.**
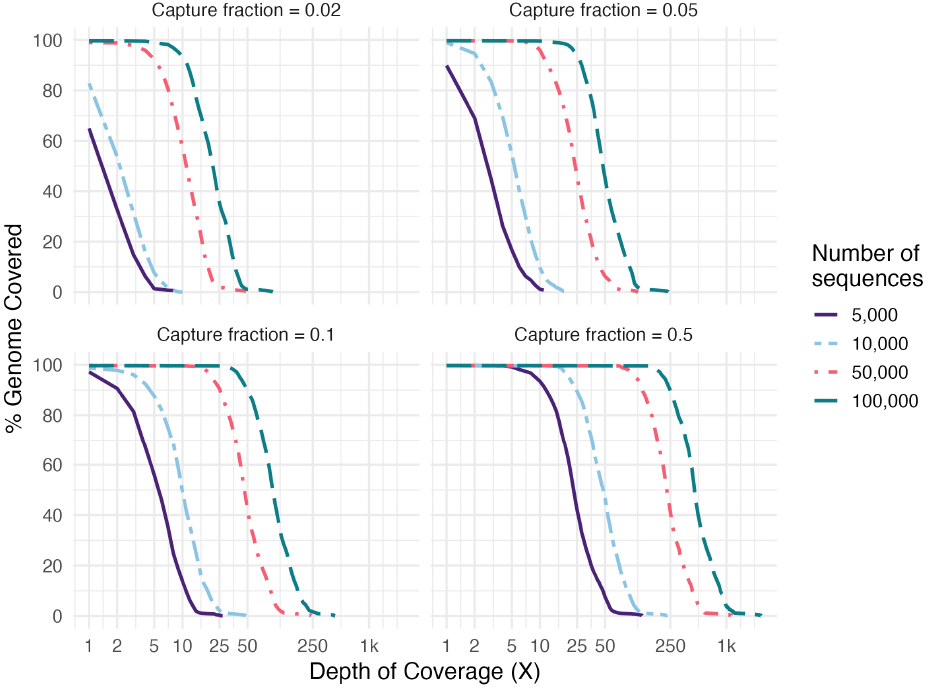
Coverage depth curves for Zika virus (ZIKV) detection against an Aedes aegypti background, sweeping capture fraction (panels) and number of simulated reads (lines, 5,000–100,000). Each curve shows the percentage of the Zika virus genome covered at or above a given sequencing depth.

These simulations allow users to identify the minimum combination of capture efficiency and sequencing depth needed for their downstream analysis goals before committing to wet-lab experiments.

## 3 Validation Using Public Data

To evaluate BaitBench simulation, we used sequence and probe data from TELSeq comprising a mock microbial community (Zymo-BIOMICS Microbial Community DNA Standard II [Log Distribution]) (Slizovskiy et al. 2022). We used BaitBench to construct an input sample of targets from TELSeq, with the reported community abundances as weights, then BaitBench used the provided probes to simulate capture and sequencing. The proportions of simulated reads for each species matched the real data very well, with a Spearman correlation of 0.916, *p* = 3.4 × 10^−6)^, Lin’s concordance correlation coefficient on log_10_ proportions = 0.954, n = 20 species in both; see **Figure 2**.

**Figure 2.**
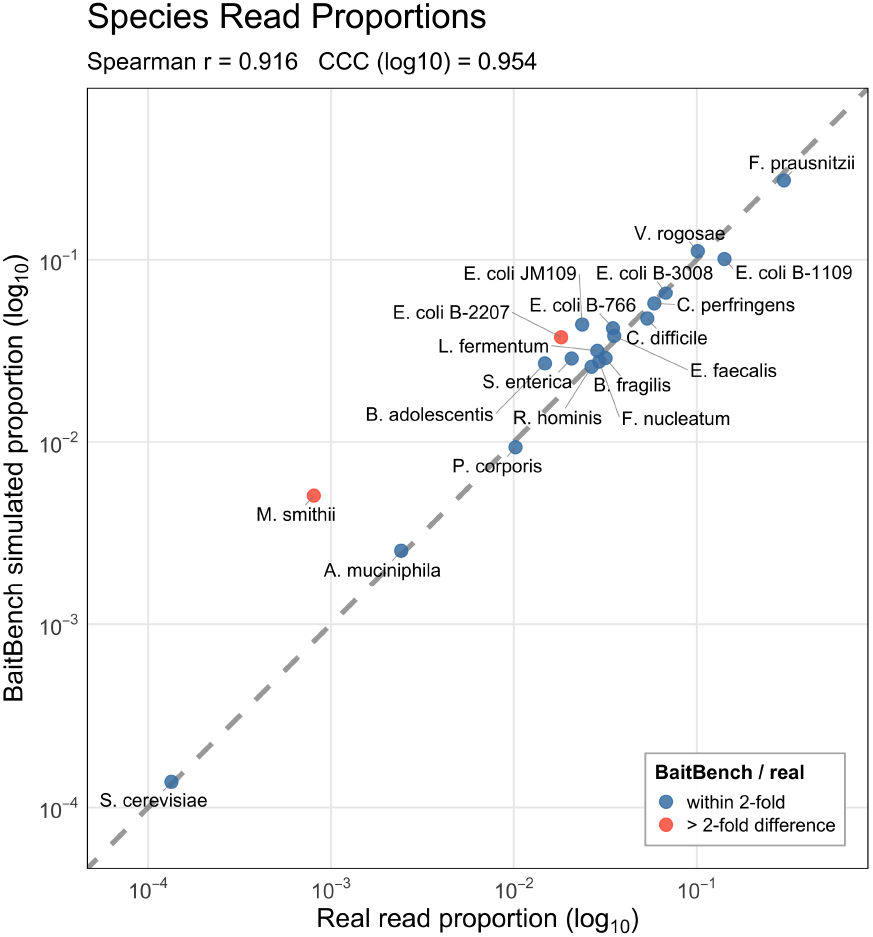
Validation of BaitBench simulations using real data. A mock microbial community of twenty species was constructed, captured, and sequenced by TELSeq (Slizovskiy et al. 2022). Using the published probe set, BaitBench simulated capture and sequencing. Agreement between simulation and experimental data is very high (Spearman’s ρ = 0.916, p = 3.4 × 10^™6^, Lin’s concordance correlation coefficient on log_10_ proportions = 0.954, n = 20 species detected in both).

Of the mismatching samples, the *M. smithii* outlier points to a limitation in our model concerning probe selection. Probes are selected uniformly from all probes that map anywhere, and only after a probe is selected are the thermodynamics considered. This procedure can over-select rare species such as *M. smithii*, with only 6 reads in the real data. The natural correction, weighting probe selection by target sequence abundance, leads to the opposite problem of over-selecting common species. Fully compensating for this would require modeling probe concentration and usage, introducing computational complexity and biological parameters users may not have access to or knowledge of. Rather than attempting to resolve these interdependencies, BaitBench uses the capture fraction parameter as a pragmatic proxy for these combined effects.

## 4 Discussion

BaitBench provides a unified tool for target capture probe analysis supporting building, quality analysis, and performance simulation in a single installation. While tools for individual steps exist, their fragmented formats and dependencies impose a significant barrier to rigorous pre-synthesis evaluation. By integrating native Rust reimplementations of CATCH and Syotti, alignment-based quality control, and thermodynamic simulation, BaitBench makes systematic probe evaluation accessible to both specialist and non-specialist users.

Because BaitBench controls all aspects of a simulation and tracks reads throughout the pipeline, it can provide a three-way classification scheme addressing a gap in existing evaluation frameworks. This allows a user to know if a call for a specific target actually originated from that target, from a different target, or from background signal. Each of these cases has different remedies: cross-reactivity may be resolved through panel redesign, while off-target capture requires stricter wash conditions or additional filtering. BaitBench’s separate False Positive target and False Positive distractor counts make this distinction explicit and actionable.

Additional use cases include assessing whether new viral strains would be detected by an existing panel, comparing commercial probe sets before purchase, or exploring how sample composition affects detection of rare pathogens in the presence of common ones. BaitBench becomes even more useful when paired with wet-lab validation. If a limit-of-detection assay is performed or known samples are run, capture efficiency can be estimated and parameters more precisely tuned. Subsequently sequencing depths may be even more accurately predicted for future experiments, or protocol change effects can be estimated. *In silico* experimentation was a major motivation for BaitBench. While actual lab work is always needed, we aim to reduce the need for trial and error at the bench, and allow more informed experiment design.

BaitBench was built to be as modular as possible. As new tools become available, they will be incorporated. As our own lab and others produce more data, we will refine our models. In particular we hope to be able to use PCR CT scores as input for sample weighting to give users a direct clinical metric for comparison.

The BaitBench command line interface runs on Linux and Mac, and GUIs are available for Mac and Windows. The Windows GUI will run natively on Windows machines without the need for an emulator or the Windows Subsystem for Linux. All wrapped tools that cannot run on Windows have had their dependencies rewritten in Rust and have become part of the distributed binary. BaitBench aims to be usable by all, regardless of computational savvy or operating system.

## Distribution

All code is available at github.com/niel-infante/BaitBench. Experts can clone the repository, install dependencies with Conda, and compile from source. Others can download the installers for Mac and Windows available at github.com/niel-infante/BaitBench/releases.

## Data availability

An archived version of the github repository is available at https://doi.org/10.5281/zenodo.23111259

